# Molecular Identification and Characterization of Two Rubber Dandelion Amalgaviruses

**DOI:** 10.1101/229443

**Authors:** Humberto Debat, Zinan Luo, Brian J. Iaffaldano, Xiaofeng Zhuang, Katrina Cornish

## Abstract

The *Amalgaviridae* family comprise persistent viruses that share the genome architecture of *Totiviridae* and gene evolutionary resemblance to *Partitiviridae*. Two genera have been assigned to this family, including genus *Amalgavirus* consisting in nine recognized species, corresponding to plant infecting viruses with dsRNA monosegmented genomes of ca. 3.4 kb. Here, we present the molecular characterization of two novel viruses detected in rubber dandelion (*Taraxacum kok-saghyz*). The sequenced viruses are 3,409 and 3,413 nt long, including two partially overlapping ORFs encoding a putative coat protein and an RNA-dependent RNA polymerase (RdRP). Phylogenetic insights based on the RdRP suggest them to be members of two new species within the *Amalgavirus* genus. Multiple independent RNAseq data suggest that the identified viruses have a dynamic distribution and low relative RNA levels in infected plants. Virus presence was not associated with any apparent symptoms on the plant hosts. We propose the names rubber dandelion latent virus 1 & 2 to the detected amalgaviruses; the first viruses to be associated to this emergent and sustainable natural rubber crop.

## Introduction

Natural rubber is an essential material to the manufacture of 50,000 different rubber and latex products. A steadily increasing demand cannot be met only by the industrial exploitation of the rubber tree (*Hevea brasiliensis*). Viable alternative crops that could be established may supplement the demand, with carbon footprint savings, which is currently supported by diverse synthetic rubbers [1] (Cornish, 2017). Rubber dandelion (*Taraxacum kok-saghyz*) is currently being developed as a sustainable source of natural rubber. Thus, a robust metabolomic, genomic, and transcriptomic characterization should advance in parallel to explore the biological landscape of this important natural resource [2] (Zhang *et al*., 2017). An additional aspect of this process is the exploration of potential microbes that could be associated with rubber dandelion. Regarding viruses, there are studies reporting the incidence of a virus in the related common dandelion (*T. officinale*), the dandelion yellow mosaic virus (*Secoviridae; Sequivirus –* [3] Bos *et al*., 1983). In addition, *T. officinale* has been described to be a reservoir host of important viral disease agents such as tomato ringspot virus (*Secoviridae; Nepovirus –* [4] Mountain *et al*., 1983), and tomato spotted wilt orthotospovirus (*Tospoviridae; Orthotospovirus –* [5] Groves *et al*., 2002). Interestingly, there are no reports describing viruses associated with rubber dandelion. Several members of *Amalgaviridae*, a relatively new family of plant and fungal viruses have been identified in the last decade ([6] Sabanadzovic *et al*., 2009). Amalgaviruses are conformed by a monosegmented dsRNA genome, containing two overlapping ORFs and a common evolutionary history. Amalgaviruses are persistent, appear to be cryptic, share the genome architecture of members of the *Totiviridae* family and gene evolutionary resemblance to partitiviruses [7] (Martin *et al*., 2011). In this report, we present the characterization of two novel viruses, tentative members of the *Amalgaviridae* family and the *Amalgavirus* genus: the first viruses associated with rubber dandelion.

## Materials and Methods

The first *T. kok-saghyz* transcriptome and associated reads as described by (Luo *et al*., 2017) were used as input for virus discovery. This transcriptome was produced from total RNA extracted from of six month old root samples of *T. kok-saghyz*, of individuals of six different genotypes characterized by dissimilar rubber yields at The Ohio State University. RNA samples were sequenced by Illumina Hiseq2000, obtaining 357,694,286 paired-end reads 100bp reads (NCBI Short Read Library (SRA) accession numbers: genotype TK6 (SRR5181667); TK9 (SRR5181665); TK10 (SRR5181664); TK14 (SRR5181663); TK18 (SRR5181662); TK21 (SRR5181661)). The sequenced reads were quality evaluated using the FASTX-Toolkit, with a cut-off score of 30 (-q). The filtered reads then went through Trinity *de novo* assembly (version 2.2.0) using standard parameters. Further, the filtered and normalized assembly (NCBI Transcriptome Shotgun Assembly (TSA) accession number GFJE00000000) of 55,532 transcripts was assessed by bulk searches using as query the complete virus refseq database available at ftp://ftp.ncbi.nlm.nih.gov/refseq/release/viral/ in a local server. BLASTX with an expected value of 10e-5 was employed as threshold, and hits were explored by hand. Tentative virus contigs were curated by iterative mapping of reads using Bowtie2 http://bowtie-bio.sourceforge.net/bowtie2/index.shtml virus Fragments Per Kilobase of transcript per Million mapped reads (FPKM) were estimated with Cufflinks 2.2.1 http://cole-trapnell-lab.github.io/cufflinks/releases/v2.2.1/. Virus annotation was implemented as described elsewhere [8–9] (Debat and Bejerman, 2019; Debat et al, 2019). In brief ORFs were predicted by ORFfinder https://www.ncbi.nlm.nih.gov/orffinder/ translated gene products were assessed by InterPro https://www.ebi.ac.uk/interpro/search/sequence-search and NCBI Conserved domain database v3.16 https://www.ncbi.nlm.nih.gov/Structure/cdd/wrpsb.cgi to predict domain presence and architecture. The 3D structure of the putative coat proteins was determined with the EMBOSS 6.5.7 Tool Garnier http://www.bioinformatics.nl/cgi-bin/emboss/garnier and coiled coil regions were predicted with COILS https://embnet.vital-it.ch/software/COILS_form.html using a MTIDK matrix. Predicted protein similarity plots were generated with Circoletto http://tools.bat.infspire.org/circoletto/ setting as E-value 10e-10. Phylogenetic insights based in predicted virus proteins were generated by MAFTT 7 https://mafft.cbrc.jp/alignment/software/ multiple amino acid alignments and FastTree 2.1.5 maximum likelihood phylogenetic trees computing local support values with the Shimodaira-Hasegawa test http://www.microbesonline.org/fasttree/. The FreeBayes v0.9.18S tool with standard parameters was employed for SNPs prediction https://github.com/ekg/freebayes. Results were integrated and visualized in the Geneious 8.1.9 platform (Biomatters, inc.).

## Results

The first publically-available RNA-Seq based *T. kok-saghyz* transcriptome, which was developed from pools of roots of genotypes with high and low rubber yields [10] (Luo *et al*., 2017), was subjected to bulk BLASTX-NCBI searches using as query the complete virus refseq database. Interestingly, two transcripts presented consistent sequence identity to the amalgavirus southern tomato virus [6] (Sabanadzovic *et al*., 2009) (50% identity at the aa level; E-value = 0.0) and blueberry latent virus (49% identity at the aa level; E-value = 0.0). The corresponding transcripts were polished by iterative mapping of RNA reads, which gave a mean coverage support of 49.1 X and 77.7 X, respectively. The curated 3,409 nt and 3,413 nt long sequences were further explored in detail and designated tentatively isolate OH of rubber dandelion latent virus 1 & 2 (RdLV1 & RdLV2).

The determined virus sequence of RdLV1 contains a 143 nt 5′UTR, a 97 nt 3′UTR, and two partially overlapping ORFs on the positive strand (Figure 1.A). The predicted ORF1 encodes a 387 aa putative Coat protein (CP) based on the first 10 BLASTP hits all corresponding to plant amalgaviruses CPs (E-value 8e-20 to 1e-78). The overlapping ORF2 encodes an 825 aa RdRP with a corresponding RNA_dep_RNAP domain (Pfam: pfam00680, E-value = 1.30e-07) at the 360-544 aa coordinates. Genome position 981 (_$) presents a putative “slippery” sequence of the form ACU_UUU_CGC suggesting a host ribosomal +1 frameshift signal that could induce the generation of a 1,055 aa, 120 kDa fusion protein (Figure 1.B). This slippery sequence is identical to the reported frameshifting signal of the amalgavirus rhododendron virus A [11] (Sabanadzovic *et al*., 2010). In addition, the RdLV2 virus sequence presents a 171 nt 5′UTR and a 100 nt 3′UTR (Figure 1.A). The predicted ORF1 encodes a 377 aa putative CP, sharing a 21.5% aa pairwise identity with RdLV1 CP. The overlapping ORF2 encodes a 749 aa RdRP with a RNA_dep_RNAP domain (pfam00680, E-value = 2.99e-10) at the 304-473 aa coordinates. Genome position 946 (_$) presents a putative “slippery” sequence CAG_UUU_CGU that could induce the generation of a 1,046 aa, 118 kDa fusion protein (Figure 1.B). The UTR regions of RdLV1 & RdLV2 are relatively A+U rich, as described for amalgaviruses [6] (Sabanadzovic *et al*., 2009), ranging from 53.1 % in the RdLV1 5′UTR to 61% in the 3′UTR of RdLV2. The putative CP of RdLV1 & RdLV2 were subjected to 3D structure prediction with the EMBOSS 6.5.7 Tool Garnier and coiled coil determination by COILS with a MTIDK matrix. A comparison of these predictions to that of representative members of *Amalgavirus* members (Figure 1.C) suggests that RdLV1 & RdLV2 present a typical α-helical central region with high probability of coiled coil as part of its tertiary structure, as is prevalent in members of family *Amalgaviridae* [12] (Nibert *et al*., 2016). It is worth noting that the predicted sequences of potential slippery sequences of RdLV1 & RdLV2 are of the general form UUU_CGN, similar to the experimentally validated sequence of influenza A virus [13] (Firth *et al*., 2012). Hypothetically, the ribosome may stall on a slippery sequence, making a pause at a rare codon (such as CGN = R) for which scarce tRNAs might be available. This pause may lead to a movement forward of one nucleotide. Translation resolves on the advanced ribosome in the +1 frame (Figure 1.B). This phenomenon has been predicted to be widespread among plant amalgaviruses [12] (Nibert *et al*., 2016). Sequence comparisons indicate that RdLV1 & RdLV2 share a 55.9% genome nt identity and a 49.5% aa pairwise identity between their predicted RdRPs. Their proposed assignment as separate species is consistent with the species demarcation criteria for the genus *Amalgavirus* proposed by the International Committee on Taxonomy of Viruses (ICTV), which specifies an amino acid sequence divergence of over 25% at the RdRPs [14]. The structural highlights of RdLV1 & RdLV2 were compared to that of ICTV recognized amalgavirus species (Table 1). The predicted genome lengths and architectures, ORFs, UTRs, gene products, protein sizes, and general viral sequence cues are consistent with the proposed assignment of RdLV1 & RdLV2 viruses as member of species linked to the *Amalgavirus* genus. The predicted RdRP of RdLV1 & RdLV2 were employed to assess evolutionary insights of the identified viruses. Maximum-likelihood phylogenetic trees of RdLV1 & RdLV2, and reported amalgaviruses, in the context of related viral families were generated based on MAFTT protein alignments. The resulting trees evidently place RdLV1 & RdLV2 in a clade of plant amalgaviruses, distant from viruses members of the *Partitiviridae* and *Totiviridae* families (Figure 2.A). RdLV1 clusters with allium cepa amalgavirus 1-2 [12] (nibertx *et al*., 2016), and RdLV2 with spinach amalgavirus 1 [15] (Park & Hahn, 2017). The complete fusion protein (FP) of RdLV1 & RdLV2 was explored in sequence similarity among representative amalgaviruses and distantly related species (Figure 2.B) using the Circoletto tool [16] (Darzentas, 2010), highlighting a robust link among the FP of RdLV1 & RdLV2 and reported amalgaviruses. Interestingly, sequence identity significantly falls beyond the *Amalgavirus* genus. Further, similarity with a species member of a new genus *Zybavirus* of fungi derived *Amalgaviridae*, the Z*ygosaccharomyces bailii virus Z* [14,17] Depierreux *et al*., 2016; Sabanadzovic et al., 2018), is consistently low (Table 1), supporting that both RdLV1 & RdLV2 could be members of the *Amalgavirus* genus of plant infecting viruses.

**Figure 1.**
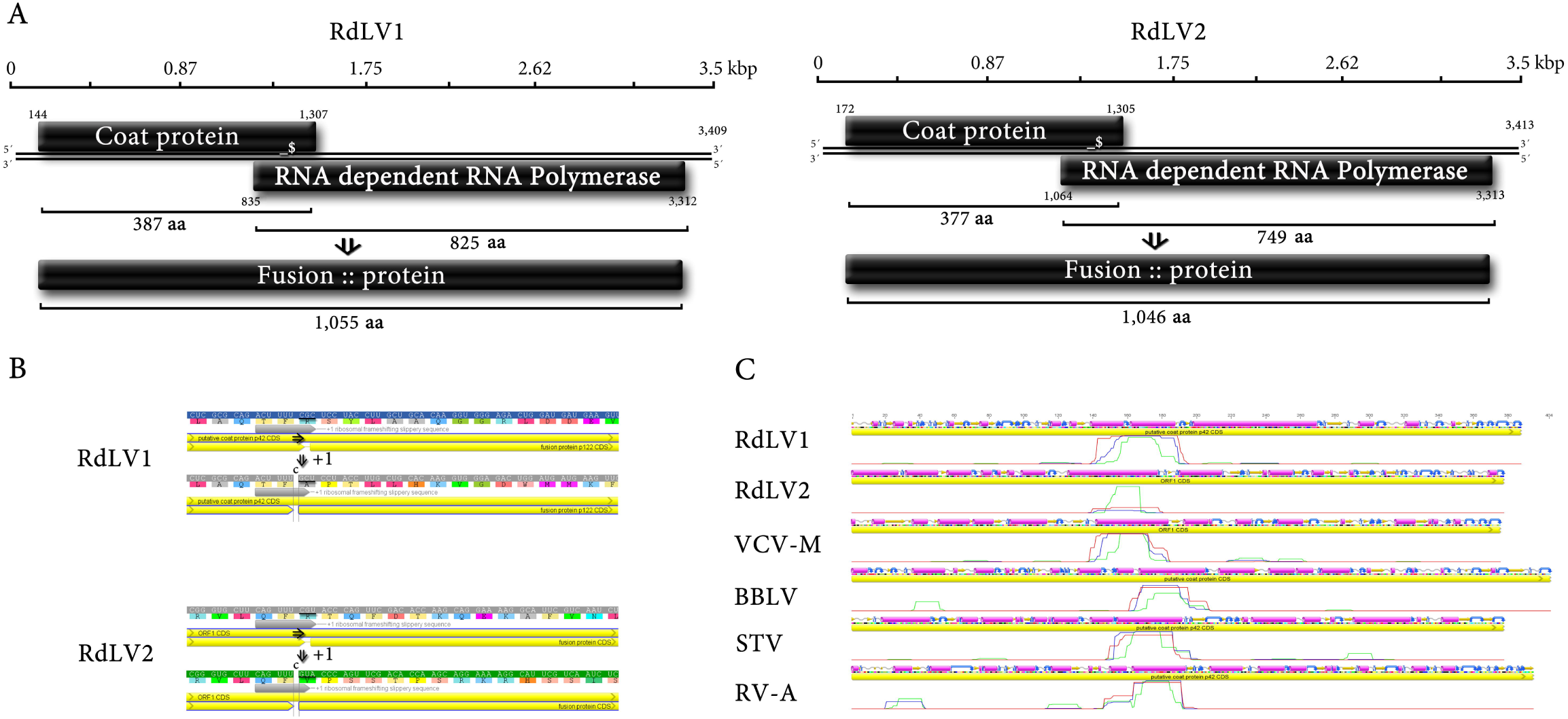
Molecular characterization of RdLV1 & RdLV2 (**A**) Rubber dandelion latent virus 1 & 2 (RdLV1 & RdLV2) linear monopartite dsRNA sequences are 3,409 and 3,413 nt long, arranging a translation strategy based in two partially overlapping ORFs. The RdLV1 virus sequence presents a 143 nt 5′UTR and a 97 nt 3′UTR. The predicted ORF1 encodes a 387 aa putative Coat protein. The overlapping ORF2 encodes a 825 aa RNA dependent RNA Polymerase. Genome position 981 (_$) presents a putative “slippery” sequence that could induce the generation of a 120 kDa fusion protein (FP). The RdLV2 virus sequence presents a 171 nt 5′UTR and a 97 nt 3′UTR. The predicted ORF1 encodes a 377 aa putative Coat protein. The overlapping ORF2 encodes a 749 aa RdRP. Genome position 946 (_$) presents a putative “slippery” sequence that could induce the generation of a 118 kDa FP. (**B**) Potential programmed ribosomal frameshifting of RdLV1 & 2. The RdLV1 ACU_UUU_CGC motif and RdLV2 CAG_UUU_CGU motif, of the general form UUU_CGN, are +1 ribosomal frameshifting motif prevalent among most plant amalgaviruses. (**C**) 3D structure prediction of the corresponding Coat proteins of RdLV1 & 2 and of representative amalgaviruses, assessed with the EMBOSS 6.5.7 tool Garnier represented on top, and coiled coil determination by COILS with a MTIDK matrix as a line graphs. Regions of high coiled-coil probability are constrained to the typical α-helical central region of the amalgavirus CPs.

**Figure 2.**
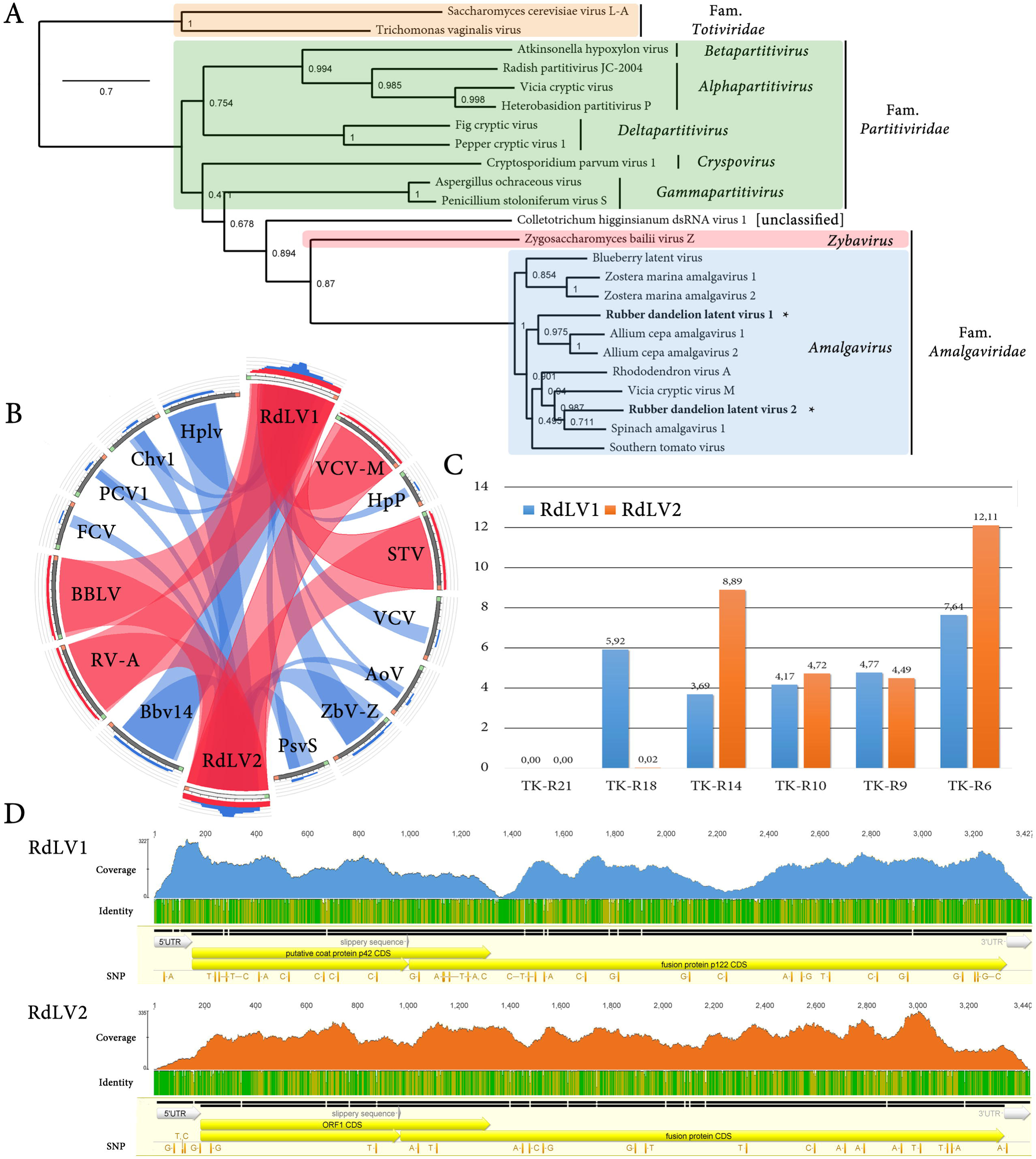
Phylogenetic insights, exploratory prevalence and RNA levels of RdLV1 & RdLV2 (**A**) Maximum-likelihood phylogenic tree of the RdRP predicted protein of reported amalgaviruses in the context of related viral families based on a MAFTT multiple alignments. Numbers at the nodes indicate percentage of bootstrap consensus support values obtained for 1,000 replicates. (**B**) Sequence similarity levels of amalgaviruses RdRP proteins among the *Amalgavirus* genus and of related viruses expressed as Circoletto diagrams. RdRPs are depicted clockwise, and sequence similarity is visualized from blue to red ribbons representing low-to-high sequence identity. (**C**) Virus RNA levels expressed as FPKM of NGS sequenced rubber dandelion total RNA root samples. Values for RdLV1 are depicted in blue columns and values for RdLV2 in orange columns. (**D**) RNAseq based read mapping graphs of RdLV1 and RdLV2 with the six combined RNA libraries. Tracks from top to bottom represent coverage per base, sequence identity from red to green (higher), and SNP prediction. GenBank accession numbers and abbreviations for the respective viruses are southern tomato virus (STV, NC_011591), rhododendron virus A (RV-A, NC_014481), blueberry latent virus (BBLV, NC_014593), vicia cryptic virus M (VCV-M, EU371896), hubei partiti-like virus 59 (Hplv, APG78262), beihai barnacle virus 14 (Bbv14, APG78182), *Zygosaccharomyces bailii* virus Z (ZbV-Z, KU200450), *Colletotrichum higginsianum* dsRNA virus 1 (Chv1, NC_028242), heterobasidion partitivirus P (HpP, AAK52739), radish partitivirus (AY748911), vicia cryptic virus (VCV, EF173396), *Saccharomyces cerevisiae* virus L-A (ScV-LA, NC_003745), *Penicillium stoloniferum* virus S (PsvS, NC_007539), *Aspergillus ochraceous* virus (AoV, EU118277), *Cryptosporidium parvum* virus 1 (CpV1, CPU95995), pepper cryptic virus 1 (PCV1, JN117276), *Trichomonas vaginalis* virus (TvV, NC_003824), fig cryptic virus (FCV, NC_015494), *Atkinsonella hypoxylon* virus (NP_604475).

To confirm the presence of the identified viruses and explore their preliminary prevalence, we further investigated six independent root total RNA samples of *T. kok-saghyz*, which were individually sequenced by Illumina Hiseq2000 generating over 291 million 100 bp pair end reads, ranging between 5.2 Gb to 6.7 Gb per sample. Interestingly, the presence of the cognate viruses was confirmed in five of the six samples by mapping of sequencing reads to the reference transcripts of RdLV1 & RdLV2 (Figure 2.C). Virus relative RNA levels varied among samples, ranging from 3.69 FPKM for RdLV1 in TK-R14, to 12.11 FPKM for RdLV2 in TK-R6. In addition, in the TK-R18 sample, only RdLV2 was found, and both viruses were absent in TK-R21, suggesting that RdLV presence is dynamic and that mixed infections are common. *De novo* assembly of the raw RNA data and further identification of RdLV isolates on the diverse samples were carried out in order to address preliminary virus diversity. Sequence variants among samples were reduced, presenting a high degree of homogeneity. Overall identity among individuals ranged from 98.3% to 99.4%, which was roughly equivalent to the observed intra-individual identity, which ranged between 99.2% and 99.5%. A consistent identity among isolates was reported for *Blueberry latent virus*, when 35 diverse cultivars were assessed and over 99% sequence identity among isolates was observed [7] (Martin *et al*., 2011). Additionally, SNP were predicted by implementing the FreeBayes tool [18] (Garrinson & Marth, 2012), and 259 well supported variants were identified among the CDS of RdLV1 & RdLV2 (Figure 2.D). Notably, 78.37% of the polymorphisms involved the 3^rd^ position of the predicted codon, suggesting a tentative constraint to avoid amino acid changes and thus maintain structure and functional domains of the respective viruses.

## Discussion

Recurrent attempts to transmit amalgavirus via grafting and mechanical inoculation have failed. In addition, amalgavirus are very efficiently transmitted vertically via seed (70–90%), and have been associated with symptomless infections in their respective hosts (Sabanadzovic *et al*., 2010). The latter is consistent with our observations on tested rubber dandelions, which could not be linked with symptoms or altered phenotypes. Future studies should explore whether RdLV1 & RdLV2 share the biological properties of persistence and exclude potential horizontal transmission. To our knowledge, there are no reports of interspecific transmission of amalgaviruses, and transmission by potential vectors has not been conclusively ruled out. Even though there are only ten species of amalgaviruses recognized by the ICTV [14] (Sabades et al 2019), recent reports suggest that the diversity of this family of viruses is much more complex and widespread among plants [12] (Nibert *et al*., 2016). Notably, the discovery of potentially cryptic viruses has been hampered by the targeted study of symptomatic organisms, which lead to the biased discovery of pathogenic viruses [19] (Geoghegan & Holmes, 2017). Next generation sequencing is unraveling a new multifaceted virosphere paradigm, where viruses are widespread and associated to every organism [20] (Greninger, 2017). The identified RdLV1 & RdLV2 correspond to the first viruses associated with *T. kok-saghyz*. RdLV1 & RdLV2 present the genomic architecture and structural highlights of amalgaviruses, which is also supported by phylogenetic insights. Our results suggest that RdLV1 & RdLV2 appear to be prevalent, commonly found in mixed infections and at low relative RNA levels and potentially cryptic. The molecular characterization of these prospective members of the *Amalgaviridae* family is a first step on the path to advance the understanding of the intriguing biology of these potential endophytes and their economically important plant host.

## Nucleotide sequences

The virus sequences of Rubber dandelion latent virus 1 & 2 have been deposited in NCBI GenBank under accession numbers MF197380 and MF197379, respectively.

## Author contributions

HD and KC conceived the study, ZL, BJI and XZ conducted the experiments, HD, ZL, BJI and XZ analyzed the data, HD wrote the manuscript. All authors reviewed and approved the manuscript.

## Compliance with ethical standards

### Research involving human or animal participants

This article does not contain any research involving human or animal participants.

---

**Table 1** Diverse structural highlights of RdLV 1 & 2 in comparison with members of ICTV recognized species of family *Amalgaviridae*. Abbreviations: Gen: virus genus, A: *Amalgavirus*, Z: *Zybavirus*, GS: Genome size (nt), 5′U: 5′UTR length (nt), OR1: ORF 1 length (nt), OR2: ORF 2 length (nt), 3′U: 3′UTR length (nt), SLPs: Slippery sequence, SLPp: SLP position, FP: Fusion protein length (aa), RP: RdRP (aa), CP: Putative Coat protein length (aa), RPi, CPi, GSi: RdRP (aa), CP (aa), and complete genome sequence (nt) identity of the corresponding amalgavirus in relation to RdLV1.

