## Supplementary material for "Molecular Identification and Characterization of Two Rubber Dandelion Amalgaviruses": Table 1

| ***Amalgavirus*** | **Gen** | **GS** | **5´U** | **OR1** | **OR2** | **3´U** | **SLPs** | **SLPp** | **FP** | **RP** | **CP** | **RPi** | **CPi** | **GSi** | **Accession n.** |
| --- | --- | --- | --- | --- | --- | --- | --- | --- | --- | --- | --- | --- | --- | --- | --- |
| Rubber dandelion latent virus 1 | A | 3,409 | 143 | 1,164 | 2,478 | 97 | ACU_UUU_CGC | 987 | 1,055 | 825 | 387 | na | na | na | MF197380 |
| Rubber dandelion latent virus 2 | A | 3,413 | 171 | 1,134 | 2,250 | 100 | CAG_UUU_CGU | 952 | 1,046 | 749 | 377 | 49.5 | 21.5 | 55.9 | MF197379 |
| ***Southern tomato virus*** | A | 3,437 | 137 | 1,134 | 2,289 | 110 | CUU_AGG_CGU | 984 | 1,063 | 763 | 378 | 49.5 | 22.0 | 55.7 | NC_011591 |
| ***Blueberry latent virus*** | A | 3,431 | 166 | 1,128 | 2,397 | 99 | UCU_UUU_CGU | 980 | 1,055 | 799 | 376 | 46.2 | 19.5 | 54.8 | NC_014593 |
| ***Rhododendron virus A*** | A | 3,427 | 94 | 1,215 | 2,424 | 47 | ACU_UUU_CGC | 1,181 | 1,078 | 808 | 405 | 48.0 | 22.1 | 54.4 | NC_014481 |
| ***Vicia cryptic virus M*** | A | 3,434 | 142 | 1,185 | 2,277 | 117 | ACU_UUU_CGU | 986 | 1,058 | 759 | 395 | 47.3 | 19.7 | 54.8 | EU371896 |
| ***Allium cepa amalgavirus 1*** | A | 3,453 | 148 | 1,176 | 2,349 | 130 | GAG_UUU_CGU | 983 | 1,057 | 782 | 391 | 51.0 | 24.5 | 56.4 | BK010347 |
| ***Allium cepa amalgavirus 2*** | A | 3,453 | 148 | 1,321 | 2,208 | 106 | GAG_UUU_CGU | 983 | 1,065 | 735 | 390 | 53.6 | 21.6 | 56.8 | BK010348 |
| ***Spinach amalgavirus 1*** | A | 3,420 | 145 | 1,164 | 2,340 | 112 | UUC_UUU_CGG | 956 | 1,053 | 779 | 387 | 48.6 | 19.5 | 57.6 | KY695011 |
| ***Zoostera marina amalgavirus 1*** | A | 3,383 | 148 | 1,149 | 2,313 | 84 | GGU_UUU_CGU | 941 | 1,049 | 770 | 382 | 45.1 | 17.9 | 56.1 | KY783316 |
| ***Zoostera marina amalgavirus 2*** | A | 3,316 | 110 | 1,188 | 2,220 | 16 | GGU_UUU_CGU | 942 | 1,062 | 739 | 395 | 47.7 | 20.0 | 55.6 | KY783317 |
| ***Zygosaccharomyces bailii virus Z*** | Z | 3,160 | 46 | 873 | 1,980 | 77 | CUU_UUU_CGA | 911 | 1,012 | 659 | 290 | 15.8 | 12.0 | 38.1 | KU200450 |

**Molecular Identification and Characterization of Two Rubber Dandelion Amalgaviruses**

Humberto Debat, Zinan Luo, Brian J. Iaffaldano, Xiaofeng Zhuang and Katrina Cornish

**Table 1** Diverse structural highlights of RdLV 1 & 2 in comparison with members of ICTV recognized species of family *Amalgaviridae*. Abbreviations: Gen: virus genus, A: *Amalgavirus*, Z: *Zybavirus*, GS: Genome size (nt), 5´U: 5´UTR length (nt), OR1: ORF 1 length (nt), OR2: ORF 2 length (nt), 3´U: 3´UTR length (nt), SLPs: Slippery sequence, SLPp: SLP position, FP: Fusion protein length (aa), RP: RdRP (aa), CP: Putative Coat protein length (aa), RPi, CPi, GSi: RdRP (aa), CP (aa), and complete genome sequence (nt) identity of the corresponding amalgavirus in relation to RdLV1.
